# SleepInvestigatoR: A flexible R function for analyzing scored sleep in rodents

**DOI:** 10.1101/2024.04.12.588853

**Authors:** Mackenzie C. Gamble, Benjamin R. Williams, James T. McKenna, Ryan W. Logan

## Abstract

Analyzing scored sleep is a fundamental prerequisite to understanding how sleep changes between health and disease. Classically, this is accomplished by manually calculating various measures (e.g., percent of non-rapid eye movement sleep) from a collection of scored sleep files. This process can be tedious and error prone especially when studies include a large number of animals or involve long recording sessions. To address this issue, we present SleepInvestigatoR, a versatile tool that can quickly organize and analyze multiple scored sleep files into a single output. The function is written in the open-source statistical language R and has a total of 25 parameters that can be set to match a wide variety of experimenter needs. SleepInvestigatoR delivers a total of 22 unique measures of sleep, including all measures commonly reported in the rodent literature. A simple plotting function is also provided to quickly graph and visualize the scored data. All code is designed to be implemented with little formal coding knowledge and step-by-step instructions are provided on the corresponding GitHub page. Overall, SleepInvestigatoR provides the sleep researcher a critical tool to increase efficiency, interpretation, and reproducibility in analyzing scored rodent sleep.

## Introduction

Sleep-wake states in animal research are traditionally partitioned into three stages based on polysomnography recordings including wake, non-rapid-eye movement sleep (NREMS), and rapid eye movement sleep (REMS). This is often referred to as sleep staging or sleep scoring and is the foundation of sleep research from humans to animals. [1] Animals, including rodents, and human electroencephalography (EEG) sleep data share many characteristics. In both species, wakefulness and REMS are denoted by low-amplitude relatively higher frequency activity with a notably strong synchronous theta rhythm prominent in REMS. Similarly, NREMS is broadly defined by high-amplitude lower frequency activity (though in humans NREMS is separated in multiple stages). [2] This makes EEG a powerful translational tool. Appropriate categorization of these stages and their subsequent analysis is essential for making accurate diagnoses of sleep disorders, and in turn is necessary for assessing the validity of various pre-clinical animal models of such conditions. Likewise, the veracity of new interventions for sleep disorders are evaluated by how they improve different measures calculated from sleep scored data across species. Beyond primary sleep disorders the use of sleep scoring is important for understanding many brain disorders and psychiatric conditions which often present with some form of sleep perturbation. [3,4] And at its most basic level, sleep scoring is essential for interrogating the basic biology of how and why we sleep.

Many advancements have been made in automated sleep scoring in both humans [5,6] and rodents, [7] with many labs adopting some degree of automation to reduce the amount of manual scoring performed. This allows for increased efficiency and time-savings, especially in rodent research where recruiting participants is not a barrier. The pre-clinical researcher benefits from the increased ability to perform a greater number of recordings over longer durations. Not surprisingly, this increases the amount of time required to organize and analyze the subsequent scored sleep data. While several companies offer proprietary software to process rodent sleep, this software is usually designed primarily for sleep scoring and the tools provided for the actual analysis are often lacking. This includes an insufficient number of measures provided, lack of versatility, poor clean data exporting options, and limited plotting capabilities. Furthermore, these options can confer a significant cost of lab resources and labor, limiting their accessibility. While some labs have internal code and processes to overcome these problems, to date no systematic, collated, open-source functions or pipelines exist. Here, we present SleepInvestigatoR, a simple, flexible R function for the analysis of scored rodent sleep data that can be executed without extensive coding experience.

## Methods

### Data Structure and Naming Requirements

SleepInvestigatoR accepts two data structures: text files from the Sirenia acquisition software (Pinnacle Technology, Inc) and comma delimited files (.csv) of scores from any system. Comma delimited files are the most robust and the recommended input. Scores are assumed to be provided in the left most column with no header unless date and time information are also provided. If data and/or time stamped data are provided, then the user can specify either the first or second column as the scores column. The function will then choose the opposing column (either first or second) as the time information column. This information must be entered in identical format and location within each file being uploaded simultaneously. Only three states are supported: wake, NREMS, and REMS (coded as 1, 2, and 3 respectively). If different scoring nomenclature was used, SleepInvestigatoR may be re-coded to the accepted nomenclature using the score value changer option as long as they represent the same three states.

For the easiest processing of files, the following naming convention is recommended for all scored sleep files: animal identifier, treatment/condition/grouping factor, animal sex, cohort, age, and then any user chosen descriptors (e.g., Date) with each named variable separated by an underscore (e.g., Mouse1_ConditionA_Female_2_3_scores). This naming convention allows for easy incorporation of pertinent stratifying variables for streamlined analyses but is not all inclusive. Separating parts of the file name by underscores is the safest option though the function should recognize any form of punctuation between each naming variable. By default, SleepInvestigatoR assumes you have both an animal identifier variable and a grouping variable. Both of these assumptions can be overridden when the function is called, in which case it will internally delineate animal IDs and supply NAs for groups. For sex, cohort, or age “NA” can be placed in its respective location of the file name and the function will then produce a column of NAs for that variable.

### Implementation

SleepinvestigatoR was written in R, [8] a free open-source statistical programming language which can easily be accesed through the integrated development environment RStudio. The main function is ~3 mb and will efficiently run in under 30 seconds on modest desktops or laptops even when a large number of files are provided.

SleepinvestigatoR is organized into the following pipeline (Figure 1). First, the user collects and prepares their scored sleep data into the proper format and places them in a user chosen folder. Following this, the user will run the main function of SleepInvestigatoR designated by the same name. This function will take all the scored files in the previously selected folder and compute all measures based on the parameters chosen before running. SleepInvestigatoR is optimized to provide easily navigatable output options that will automatically organize results by animal identifier and treatment condition (or other designated grouping factor) when the required file naming system is provided. This file naming system also allows for simultaneous organization by common covaraites including: sex, cohort, and age. The user can indicate that a time stamp is provided for each epoch which will permit zeitgebers and light on/off information to be incorporated into the output.

**Figure 1.**
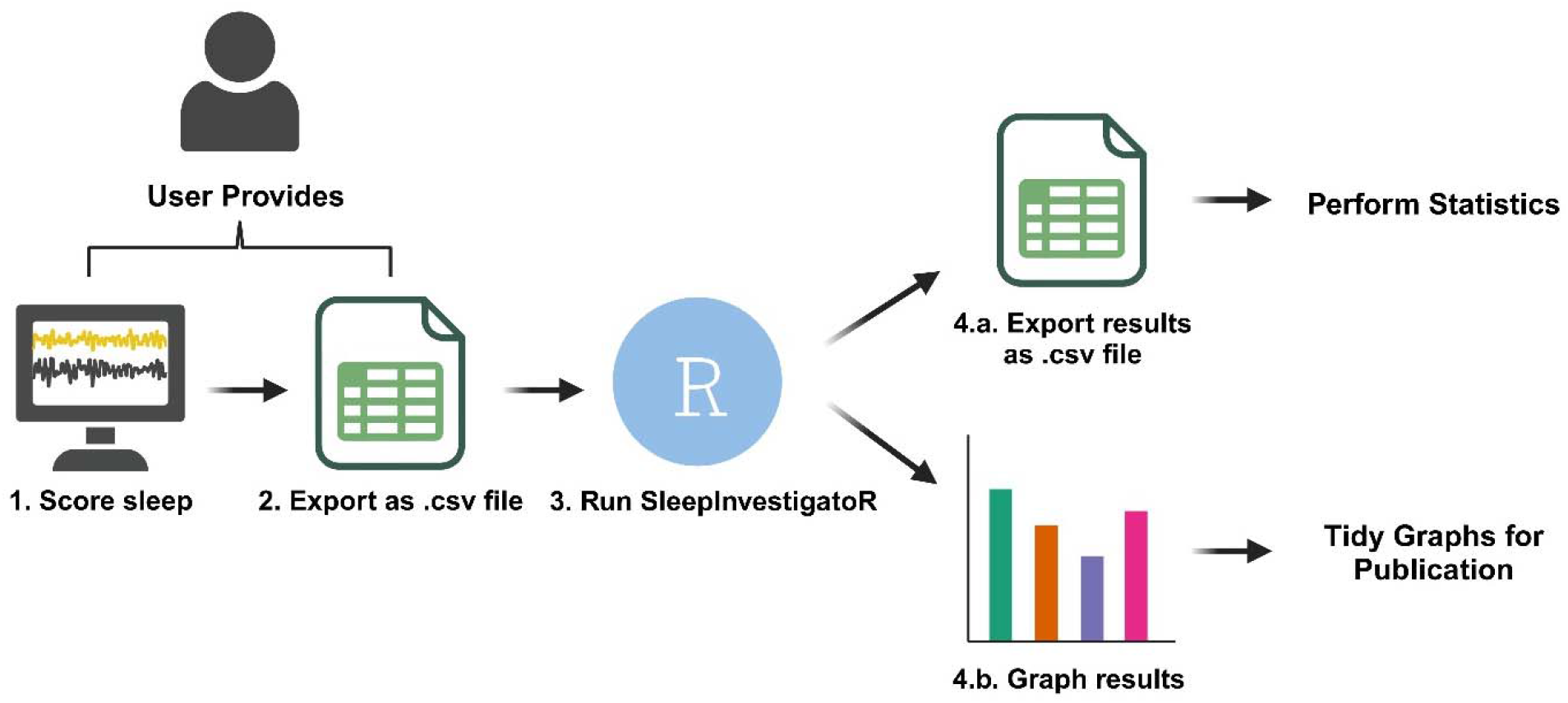
Schematic of the SleepInvestigatoR pipeline. Overview of SleepInvestigatoR pipeline. Polysomnography files are scored and exported to a central folder. SleepInvestigatoR then collates and analyzes all scored files in the designated folder. SleepInvestigatoR will then exports a summary file of measures which can also be graphed using an accessory function.

A set of 22 unique measures, or 90 total variables, are provided in long or wide format for convienent use in various statistical programs. The output includes the most common measures reported in animal studies as well human sleep research to maxmize translational value and applicability to a wide audience [9,10]. If baseline data are collected, the function can provide all measurements normalized to this baseline as well as the unormalized, absolute measures. Furthermore, a score checker is provided which when used will mark any files and the specific epoch(s) in which Wake-REMS transitions may occur, as this is usually indicative of a scoring mistake. This feature can be turned off to accommodate studies where Wake-REMS is biologically relevant (e.g., narcolepsy research)[11]. The function prioritizes flexbility and allows the user to choose the epoch length, how the sleep scoring of each animal will be divided, and the minimum length of time of a state to classify as a bout. Finally, a small plotting function is also provided to quickly produce graphs for the most commonly reported meaures.

### Parameters

There are 25 total parameters that a user can set to change the output provided by SleepInvestigatoR (Table 1). It is important that the user reads the documentation carefully and selects the appropriate parameters to generate their preferred outcome measures. By default, SleepInvestigatoR will only provide sleep-wake measures averaged over the entire recording period and assumes the default epoch length for scoring is four seconds with a latency and bout cut off time of one minute. This is all fully customizable.

**Table 1.** Parameters of SleepInvestigatoR.

| **Parameter** | **Definition** |
| --- | --- |
| FileNames* | List containing all score files to be analyzed.<br><u>Default:</u> FileNames |
| file.type* | File type of the scored files.<br><u>Default:</u> .csv |
| epoch.size* | Epoch size in seconds that sleep-wake was scored in.<br><u>Default:</u> 4 seconds |
| max.hours* | Max number of hours to be analyzed across all files. |
| byBlocks | The number of blocks to split each scored file into.<br><u>Example:</u> If set to 24 for a 24-hour file this will produce hourly data |
| byTotal | Calculates all measures across the entire recording period.<br><u>Default:</u> TRUE |
| score.checker | Checks for any Wake-REMS transitions and stops analysis if any are present identifying the file and its location in the file.<br><u>Default:</u> TRUE<br>Note: Set to FALSE to look at Wake to REM transitions (e.g., narcolepsy studies) |
| id.factor | Indicates whether an animal identifier is present.<br><u>Default:</u> TRUE |
| group.factor | Indicates whether a grouping variable is present.<br><u>Default:</u> TRUE |
| normalization | Calculates all measures normalized to a provided baseline.<br><u>Default:</u> FALSE |
| time.stamp | Indicates whether time stamped bouts are present.<br><u>Default:</u> FALSE |
| time.format | The format of the time stamps. |
|  | <u>Default:</u> hh:mm:ss |
| date.format | The format of the dates in the time stamps.<br><u>Default:</u> m/d/y |
| date.separator | The variable used to separate date and time.<br><u>Default:</u> " |
| lights.on | Time the lights turn on provided in the same format as the time stamps. |
| lights.off | Time the lights turn off provided in the same format as the time stamps. |
| score.value.changer | Can specify what values stand for Wake, NREMS, and REMS if they were not scored in the required 1,2,3 format. |
| sleep.adjust | Truncates the file so it starts at the sleep period (e.g., NREMS onset). |
| Wake/NREMS/REMS/Sleep/Propensity.cutoff | Number of epochs to count as a bout.<br><u>Default:</u> 15 (60 sec with an epoch size of 4 sec) |
| data.format | Indicate whether the data should be presented in long or wide format.<br><u>Default:</u> long |
| save.name | Chosen name of the output file. |

List of every adjustable parameter provided by SleepInvestigatorR and its purpose. Parameters designated with asterisk need to be set by the user.

### Output

SleepInvestigatoR output is a .csv file containing all measures (Table 2) which are either collapsed or grouped by time. The output can be arranged into long or wide format depending on the data format setting specified.

**Table 2.** Explanation of the list of unique measures provided by SleepInvestigatoR.

| **Metric** | **Definition** |
| --- | --- |
| Total Hours (Hours) | Total number of hours in the file |
| Latencies (Seconds) | Amount of time to:<br>A) The first bout of each state.<br>B) The first bout of wake following the initial bout of each sleep state. |
| Onsets (Epoch number) | The epoch indicating the first instance of each state |
| Onset Bout Lengths (Seconds) | The length of time of:<br>A) The first instance (onset) of each state.<br>B) The first instance (onset) of wake following each sleep state. |
| Offsets (Epoch number) | The epoch indicating the last instance of each state |
| Offset Bout Lengths (Seconds) | The length of time of the last instance of each state |
| Transition Counts (Count) | A count of the number of transitions made |
|  | between each state |
| Epoch Counts | A count of the total number of epochs of each state |
| Seconds | A count of the total number of seconds of each state |
| Percents | The percent of the total recording time spent in each state |
| Bout Counts | A count of the number of bouts of each state |
| Arousal Count | A count of the total number of instances of wake of any length |
| Microarousal Count | A count of any instance of wake less than the user set bout length |
| Average Bout Durations | The average length of time of each bout of each state |
| Propensities | The average length of time between a wake (or sleep state) to beginning of sleep state (or wake) |
| Average Interbout Intervals | The average length of time between each bout of each state |
| Wake After Sleep Onset | The total amount of time spent in wake after sleep onset |
| Wake After Sleep Offset | The total amount of time spent in wake after sleep offset |
| Sleep Fragmentation Index (per minute) | $\text{Arousal Count} / (\text{Total Sleep Time} / 60)$ |
| Total Sleep Time | The total amount of time spent in NREMS and REMS combined |
| Sleep Efficiency | The total sleep time divided by the total recording time |
| Vigilance state counts as reported by Trachsel *et al.* 1991 | A count of the total number of bouts of each state, categorized by increasing length |

### Quality Control

All code was tested using synthetic data. To ensure accuracy, manual checking was performed on all measures by at least two individuals. All code testing was performed on Windows computers but should run on Mac and Linux computers as R is compatible with both.

### Computing Requirements

SleepInvestigatoR requires only modest computing power and can run on a laptop or desktop without the need for a graphics card. All code was tested on a Lenovo Ultrabook with the following specifications:11th Gen Intel(R) Core(TM) i5-1135G7 @ 2.40GHz with 12 GB RAM running Windows 11.

### Plotting

Beyond the main analysis function, a single plotting function for viewing analyzed data is provided with SleepInvestigatorR. The function produces a plot overview which offers a quick snapshot of the data, including percent of time spent in each state, latencies to N/REMS, transition counts, bout counts, and bout durations. This function is ideal for a cursory look at the data such as checking for errors, looking at trends, and assessing the feasibility of a pilot experiment. These functions produce graphs that can easily be turned into publishable quality figures either through direct editing of the code or by exporting them as vector files for manipulation using vector editing software (e.g., Adobe Illustrator).

### Illustrative Example

As a simple proof of concept SleepInvestigatoR was used to analyze sleep scores in a publicly available dataset from Bush and colleagues. [12,13] The experiment obtained from this dataset involved the assessment of pharmacogenetic stimulation of preoptic area GABA neurons on sleep in mice.

## Data and Code Availability

All code is freely available at https://github.com/mgamble1023/SleepInvestigatoR including detailed instructions on how to run the code.

## Results

### SleepInvestigatoR replicates the NREMS enhancement finding from the Bush et al. 2022 dataset

Using the dataset publicly provided by Bush and colleagues [12,13] we were able to recreate Figure 2.C from their paper (pharmacogenetic stimulation of POA GABAergic neurons) using SleepInvestigatoR, which found NREM enhancement following a single injection of Clozapine N-oxide (CNO) (Figure 2). Additionally, using the ancillary plotting functions, SleepInvestigatoR quickly generated a graph of the results which, with minor modifications, matched the original style of the published graph (Figure 2).

**Figure 2.**
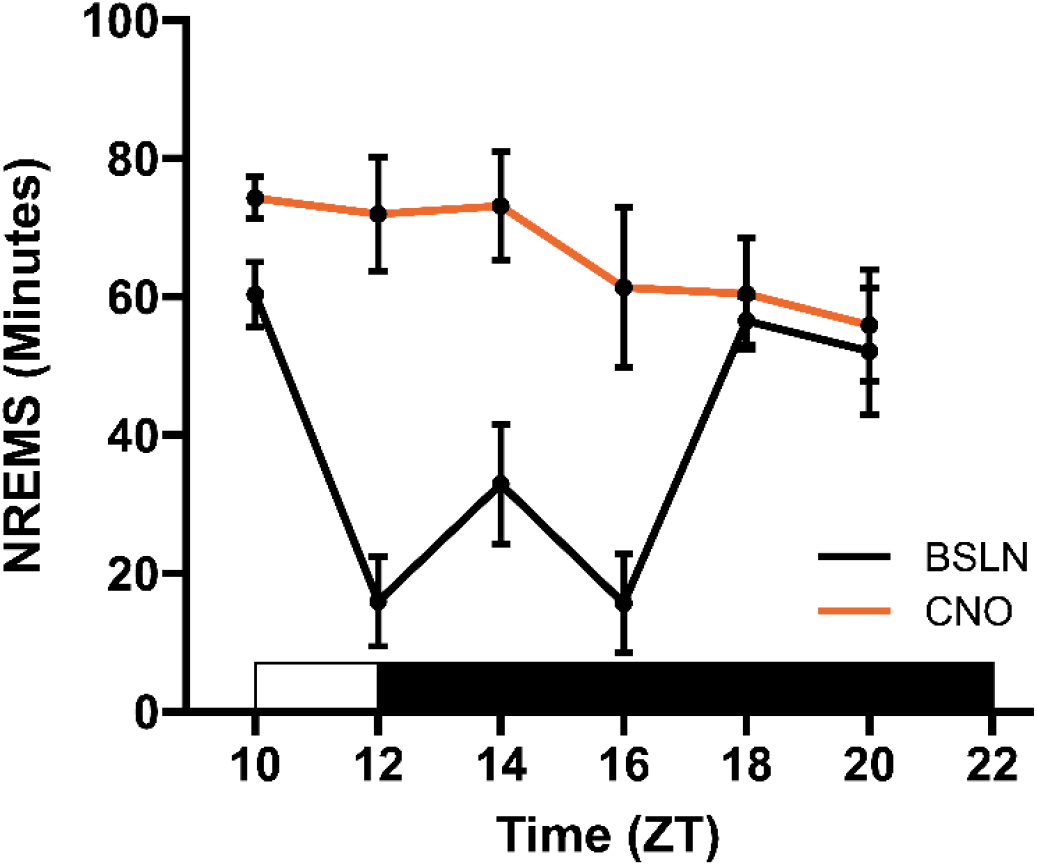
SleepInvestigatoR reports elevated NREMS following pharmacogenetic stimulation (Bush et al. dataset [10]) Replication of Figure 2. C from the Bush et al. dataset using SleepInvestigatoR. Stimulation of the preoptic area of the hypothalamus increased NREMS.

### SleepInvestigatoR reveals that POA GABA stimulation at the end of the light period increases NREMS bout count without increasing bout duration

As previously reported, cute pharmacogenetic stimulation of GABA POA neurons increased NREMS time in either the light or dark period, but only increased NREMS bout duration when administered in light period (exact ZT unspecified). [14] In the paper from Bush and colleagues CNO is administered at the end of the light cycle and changes in NREMS bout count and bout duration are not provided. Using the same dataset as before, we used SleepInvestigatoR to analyze NREMS bout count and bout duration to establish how NREMS was manipulated by pharmacogenetic stimulation. SleepInvestigatoR and subsequent statistical analysis (GraphPad Prism 10.2.2) revealed that NREMS bout count (minimum length of 60 seconds) had significant interaction effect between treatment x time (F_(5, 40)_ = 7.010, P<0.0001), but not treatment alone (F_(1, 8)_ = 2.227, P=0.1739). Post-hoc analysis indicated that the number of bouts was significantly elevated by pharmacogenetic stimulation at ZTs 12 (6.6 vs 22.0, P=0.0363) and 16 (4.4 vs 16.4, P=0.0494) (Figure 3. A.). NREMS bout duration had no significant effects due to treatment alone (F_(1, 8)_ = 0.5964, P=0.4622) or treatment x time (F_(5, 40)_ = 1.811, P=0.1326) (Figure 3. B.)

**Figure 3.**
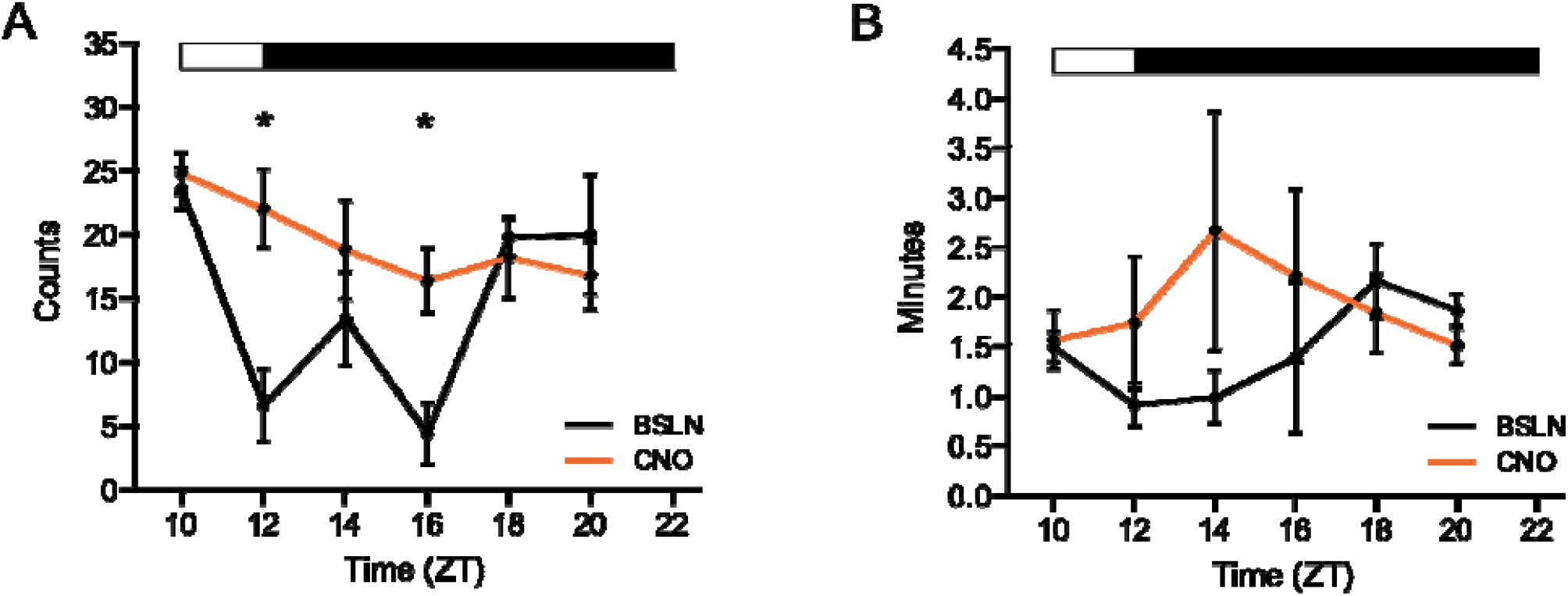
Analysis of average NREMS bout count and duration from the Bush et al. dataset using SleepInvestigatoR. Extended analysis of NREMS following pharmacogenetic stimulation of preoptic area do the hypothalamus in the Bush et al. dataset. **A** NREMS bout counts. (**B**) Average NREMS bout duration. \**p* < 0.05

## Discussion

SleepInvestigatoR is capable of analyzing sleep of multiple animals and collecting the subsequent data into a centralized output that allows for easy plotting and interpretation. Using SleepInvestigatoR is considerably faster than manual analysis and provides a comprehensive list of measures that are valuable across multiple areas of investigation within the sleep field. Measures beyond gross sleep-wake state measures may reveal more nuanced and biologically relevant changes in the sleep researcher’s experiments. As we show using the Bush and colleague’s dataset, [12,13] their pharmacogenetic treatment increased NREMS bouts, but not duration. These differences can mark important differences between various psychiatric and neurological diseases. For instance, bipolar disorder is associated with increased REMS density [15] while Huntington’s disease on the other hand is associated increased time awake after sleep onset [16]. In addition, these more subtle sleep changes can have distinct discrete behavioral impacts. For example, a study by Rolls and colleagues found that decreasing NREMS bout duration without altering overall time in NREMS via optogenetics impairs memory consolidation mice. [17] SleepInvestigatoR is able to save the researcher time and quickly calculate these measures without the need for extensive computing power or coding expertise. Moreover, SleepInvestigatoR helps contribute to reproducibility providing an open source and standardized framework for analyzing scored sleep as well as decrease the odds of human error.

Despite SleepInvestigatoR’s advantages there are several notable limitations. While animal sleep is usually just scored as wake, NREMS or REMS, it can be beneficial to make other categorizations such as mixed/in-between states, [18] light vs deep NREMS or slow wave sleep, [19,20] quiet vs active wake, [21–23] and pre-REMS. [24,25] SleepInvestigatoR is only equipped, however, for analyzing data coded as wake, NREMS, and REMS. Additionally, while SleepInvestigatoR allows manual cutoff lengths for bouts it does not allow out-of-box categorization of different bout lengths such as distinguishing between long wake vs short wake. [26] Since SleepInvestigatoR is open source, all these limitations present opportunities for future work to build and expand upon the code.

While scoring of sleep itself is the more arduous and time-consuming task, analyzing sleep stages can become increasingly tedious as the amount and length of recording increases. With many advances being made in automated sleep scoring it is becoming increasingly easy to generate more data, and in turn a new bottleneck has surfaced. Simultaneously, this also increases the odds of human error. SleepInvestigatoR was designed to overcome these obstacles and facilitate expedient analysis of scored rodent sleep. In summary, SleepInvestigatoR can increase efficiency, interpretation, and reproducibility in rodent sleep research.

## Funding

NIMH R21MH125242 and VA I01BX002774, J.T.M.; NHLBI R01HL150432 and R01HL150432-S1, R.W.L.

## Disclosure Statements

### Nonfinancial conflict of interest

J.T.M. is a Research Health Scientist at VA Boston Healthcare System, West Roxbury, MA. The contents of this work do not represent the views of the US Department of Veterans Affairs or the United States Government. J.T.M. received partial salary compensation and funding from Merck MISP (Merck Investigator Sponsored Programs) but has no conflict of interest with this work. Other authors declare no competing interests.

### Financial conflict of interest

The authors have no financial disclosure to declare.

## Notes

### Competing Interest Statement

The authors have declared no competing interest.

